# A viral transduction approach for iPSC differentiation into cortisol-producing steroidogenic cells

**DOI:** 10.1101/2025.02.13.638168

**Authors:** Laurie P. Lee-Glover, Timothy E. Shutt

**Affiliations:** Department of Biochemistry & Molecular Biology, Cumming School of Medicine, University of Calgary, Alberta, Canada; Department of Biochemistry & Molecular Biology, Cumming School of Medicine, University of Calgary, Alberta, Canada; Department of Medical Genetics, Cumming School of Medicine, University of Calgary, Alberta, Canada; Hotchkiss Brain Institute, Cumming School of Medicine, University of Calgary, Alberta, Canada; Alberta Children’s Hospital Research Institute, Cumming School of Medicine, University of Calgary, Alberta, Canada; Snyder Institute for Chronic Diseases, Cumming School of Medicine, University of Calgary, Alberta, Canada

## Abstract

Steroid hormones are important signaling molecules that are primarily produced by specialized cells. The ability to culture steroidogenic cells is critical to study this important process. While a few steroidogenic cell lines are available for study, they are typically derived from cancer patients and do not reflect genetic alterations in steroidogenic genes that cause human disease. In this regard, iPSCs are a powerful tool for modeling human disease, as patient-derived iPSCs can be differentiated into a variety of different cell types. While other approaches exist to differentiate iPSCs into steroidogenic cells, they are typically complex and time consuming. In contrast, a simple approach based on lentivirus transduction of SF1, the master regulator of steroidogenesis, can differentiate multiple types of cultured cells into a steroidogenic state. However, we show here that this approach does not work for iPSCs. To circumvent this limitation, we report a simple adaptation that first differentiates iPSCs to embryoid bodies prior of SF1 transduction. With this modified approach, we provide a straightforward cost-effective approach to differentiate iPSCs into actively steroidogenic cells.

## Introduction

The differentiation of steroidogenic cells from various precursor cell types is of interest for a variety of research and clinical questions. Disorders of steroidogenesis include congenital adrenal hyperplasia, resulting from insufficiency of the adrenocortical steroids (1), as well as hypogonadism, resulting from deficiency of reproductive hormone production (2). Further, evidence suggests a number of other conditions involve some degree of dysregulated steroidogenesis, including Parkinson’s disease (3) and Alzheimer’s disease (4,5). Recapitulation of the differentiation of steroidogenic tissues *in vitro* can be used to study steroidogenic tissue development and primary disorders (6). Differentiating patient-derived cells into steroidogenic cells can be used to study genetic effects on steroidogenesis, as well as the efficacy of different therapeutics for personalized medicine (7). Finally, the conversion of patient-derived cells to steroidogenic cells are a potential source of cell replacement therapy/regenerative cell therapy in steroid insufficiency patients (8–10).

As first demonstrated in 1997, the transcription factor SF1 is a capable of driving a steroidogenic program when expressed in cells (11). Since then, a number of protocols have been developed in the literature to convert precursor cell types into either adrenocortical-like or Leydig-like cells (7–10,12–21). Generally, these methods involve two key elements. First, the forced expression of steroidogenic factor 1 (SF1), a transcription factor that is the master regulator of steroidogenesis (22). Secondly, treatment with a cell permeable cAMP analogue such as 8-bromo-cAMP that is needed for effective activation of steroidogenic genes by SF1 (13,23). While a number of protocols converting multipotent cells such as mesenchymal stem cells (MSCs) and fibroblasts have been established (7,9,10,12–17), induced pluripotent stem cells (iPSCs), which can be reprogrammed from various cell types, are a subject of growing interest, especially in the context of disease modelling. Studies published on this cell type so far suggest that steroidogenic differentiation is a less straightforward process (8,18–21).

Ruiz-Babot *et al.* (7) recently described a novel method for simple differentiation of patient-derived cells into adrenocortical-like cells using lentivirus-mediated SF1 expression. As this method has yet to be tested in pluripotent stem cells, we have extended this approach to iPSCs. However, we note iPSCs are not amenable to simple SF1 over-expression, a limitation that we show can be overcome by first differentiating iPSCs into embryoid bodies prior to SF1 expression.

## Methods

### Cell culture

HEK293 cells were cultured in DMEM with 10% heat-inactivated fetal bovine serum. Adrenocorticocarcinoma cell lines H295R and its clonal derivative HAC15 (24) were cultured in DMEM:F12 supplemented with ITS+ (Corning, 354352) and 2.5% Nu serum (Corning, 355100). The AIW002-02 iPSC line was obtained in collaboration with C-BIG Repository (McGill University) (25), and was cultured in mTeSR Plus (Stemcell, 100-0276) and on Matrigel (Corning, 354277) coated plates.

### Lentivirus production

Lentivirus production was adapted from previously described protocols (26). In short, HEK293 cells were seeded at 2.5×10^6 cells per 10 cm dish and transfected with 6 *μ*g transfer vector, 5 *μ*g packaging vector and 2.5 *μ*g envelope vector using calcium phosphate transfection. This transfection was scaled up as needed. Packaging vector psPAX2 (Addgene plasmid # 12260) and envelope vector pMD2.G (Addgene plasmid # 12259) were gifts from Didier Trono. Transfer vector pLOC-SF1 (shared from Dr. Leonardo Guasti at Queen Mary University of London, backbone Dharmacon OHS5832) (7). The following day, media was replaced with fresh media containing 1mM sodium butyrate. Media containing virus was collected at 48 hours, 72 hours and 96 hours post transfection. Virus supernatant was passed through a 0.45 *μ*m sterile filter and stored at -80C. Lentivirus was concentrated using Lenti-X Concentrator (Takara Bio 631232) and physical titre was determined using ABM qPCR Lentivirus Titer Kit (Applied Biological Materials LV900).

### SF1 transduction and steroidogenic differentiation

Lentivirus (400 viral particles/cell as determined by ABM qPCR Lentivirus Titer Kit) was added along with 10 *μ*g/mL hexadimethrine bromide (Polybrene) (Sigma-Aldrich, 107689) to target cells. Two days later, cell culture media was supplemented with 100 *μ*M 8-bromo-cAMP (Stemcell 73604). Also at this time, cells were selected with 10 *μ*g/mL blasticidin (Invivogen, ant-bl) until untransduced control cells were completely dead, about 7 days. Cells were treated with 8-bromo-cAMP for at least 8 days before analysis, and maintained in 8-bromo-cAMP for continued culture.

### Embryoid body differentiation

Conversion of iPSCs to embryoid bodies was carried out as described previously (27). Briefly, iPSCs were clump passaged, pelleted and resuspended in mTESR Plus media with 10 *μ*M Y-27632 (Stemcell 72308). The cells were transferred to a sterile non-tissue culture treated dish (e.g. a sterile Petri dish) to encourage embryoid body formation. The next day the media was changed to DMEM:F12 with 20% Knockout Serum Replacement (KOSR) (Gibco, 10828010). Media was changed every 1-2 days as embryoid bodies were cultured for 6 days. Embryoid bodies were then collected, resuspended in DMEM:F12 with 10% heat inactivated fetal bovine serum and replated in a standard tissue culture-treated plate to culture outgrowing adherent cells.

### Microscopy

Images were taken using Olympus CKX53 microscope with Olympus LC30 camera. The program Olympus Cell Sens Entry was used for image acquisition and addition of scale bars. Brightness and contrast adjustments were performed in ImageJ. If two images were to be compared, the exposure time during acquisition and image adjustments during editing were kept consistent.

### Western blot

Cells were harvested and lysed in RIPA buffer (Thermo Scientific, 89901) with protease inhibitor cocktail (BioVision, K272-1). The soluble protein fraction was collected as supernatant after centrifugation at 14 000xg for 30 minutes. Protein concentration was determined using bicinchoninic acid (BCA) assay (Thermo Scientific, 23223 and 23224). Unless otherwise indicated, 50 *μ*g of protein was loaded per well. Samples were mixed with 1x Laemli buffer (Bio-Rad, 1610747) with 10% beta-mercaptoethanol and boiled for 5 minutes at 95 °C. Proteins were separated on SDS-containing 10 or 12% polyacrylamide gels along with Precision Plus Protein All Blue Prestained Standards (Bio-Rad, 1610373) as molecular weight markers. The gel was transferred to a PVDF membrane overnight. Blocking and antibody incubations were performed in 5% milk in TBST. Antibodies used are listed in **Table 1**. Chemiluminescence was visualized with Clarity Western ECL Substrate (Bio-Rad, 1705061) or Super Signal West Femto (Thermo Scientific, 34095).

**Table 1.**
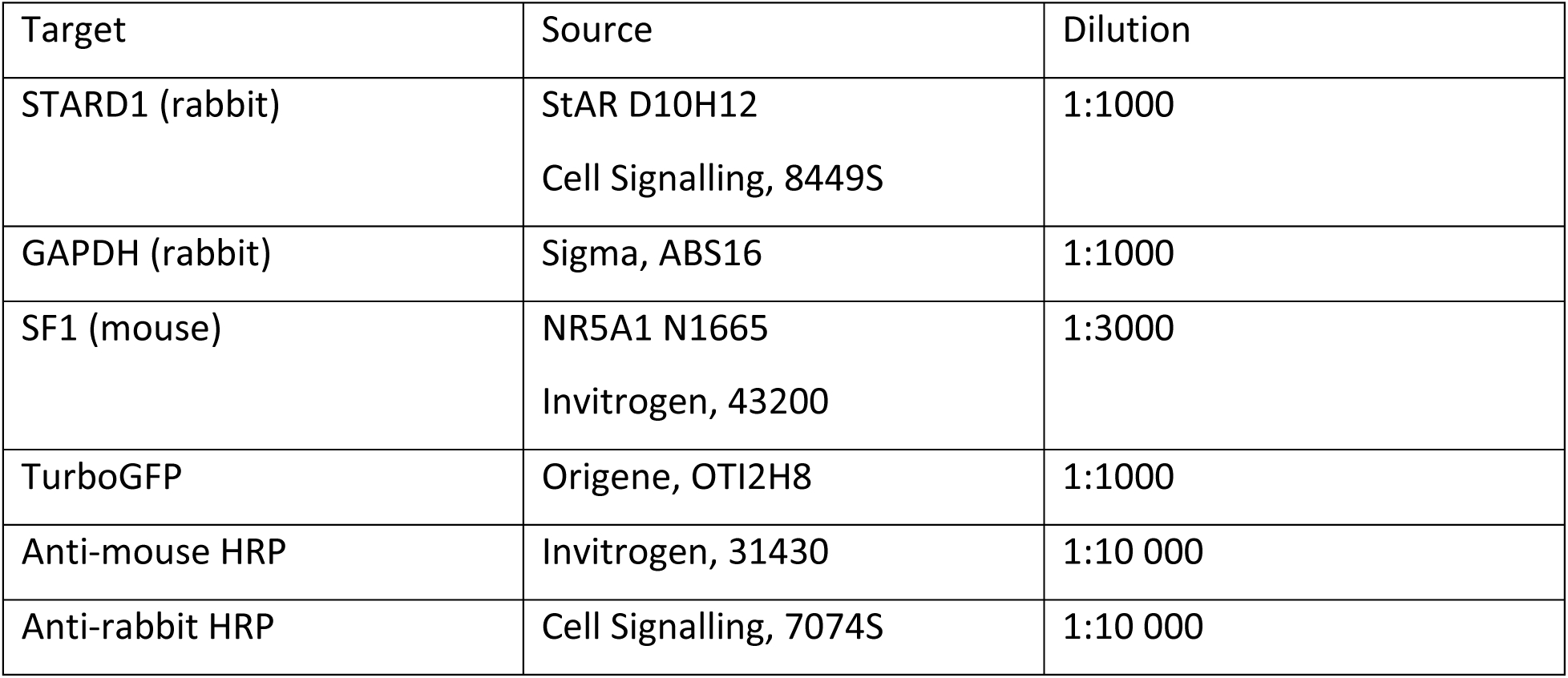

| Target | Source | Dilution |
| --- | --- | --- |
| STARD1 (rabbit) | StAR D10H12<br>Cell Signalling, 8449S | 1:1000 |
| GAPDH (rabbit) | Sigma, ABS16 | 1:1000 |
| SF1 (mouse) | NR5A1 N1665<br>Invitrogen, 43200 | 1:3000 |
| TurboGFP | Origene, OTI2H8 | 1:1000 |
| Anti-mouse HRP | Invitrogen, 31430 | 1:10 000 |
| Anti-rabbit HRP | Cell Signalling, 7074S | 1:10 000 |

### Pregnenolone and cortisol quantification

As steroid hormones are cell permeable (28), pregnenolone and cortisol were measured from cell culture media. Cells were rinsed with PBS and complete culture medium was replaced with serum-free medium supplemented with 100 *μ*M 8-Bromo-cAMP. When media was collected for pregnenolone measurement, media also included 1 *μ*M trilostane (AdooQ Bioscience, A10949-50) and 1 *μ*M abiraterone (AdooQ Bioscience, A10020-50), which inhibit 3β-hydroxysteroid dehydrogenase and 17α-hydroxylase respectively, and prevent conversion of pregnenolone to other downstream steroids. Cortisol ELISA kit (Cayman Chemicals, 500360) and pregnenolone ELISA kit (IBL America, IB59107) were used to quantify steroid concentrations following manufacturer’s instructions. In cases where pregnenolone or cortisol levels were compared between different cell lines, an additional normalization step was taken to adjust for any discrepancies in seeding densities. Cells in the well were harvested, lysed and protein concentration was quantified by BCA assay and used to normalize steroid concentrations.

### qRT-PCR

To assess gene expression, RNA was isolated from cultured cells using EZNA Total RNA Kit II (Omega Bio-Tek, R6934-01) and reverse transcribed using iScript Advanced cDNA Synthesis Kit (Bio-Rad, 1725038). PCR reactions used PowerUp SYBR Green Master Mix (Applied Biosystems, A25778) on QuantStudio6 Real-Time PCR Machine (Applied Biosystems). For analysis, the delta-delta Ct (cycle threshold) method was used using actin as an endogenous control. Primer sequences: OCT4 forwards 5’-TATCGAGAACCGAGTGAGAG-3’, OCT4 reverse 5’-TCGTTGTGCATAGTCGCT-3’, brachyury forwards 5’-TGCTGCAATCCCATGACA-3’, brachyury reverse 5’-CGTTGCTCACAGACCACA-3’, actin forwards 5’-AAGACCTGTACGCCAACACA-3’, actin reverse 5’-AGTACTTGCGCTCAGGAGGA-3’.

## Results

### Lentiviral overexpression of SF1 and steroidogenic differentiation

The pLOC-SF1 lentivirus transfer vector containing the SF1 open reading frame is reported to be able to differentiate various cell types into steroidogenic cells (7). This vector drives constitutive expression of SF1, GFP and blasticidin resistance under a single promoter (**Figure 1A**). Before attempting to differentiate iPSCs, we wanted to confirm the lentiviral transduction and steroidogenic differentiation process using HEK293 cells. After transduction and 8 days of 8-bromo-cAMP treatment, a treatment used to activate steroidogenesis (7), we observed GFP positive cells (**Figure 1B**), expression of both SF1 and STARD1 (a key protein mediating the rate limiting step of steroidogenesis) (**Figure 1C)**, as well as production of pregnenolone, the precursor to all other steroid hormones (**Figure 1D**). These results indicate that forced SF1 expression could induce steroidogenesis in HEK293 cells.

**Figure 1:**
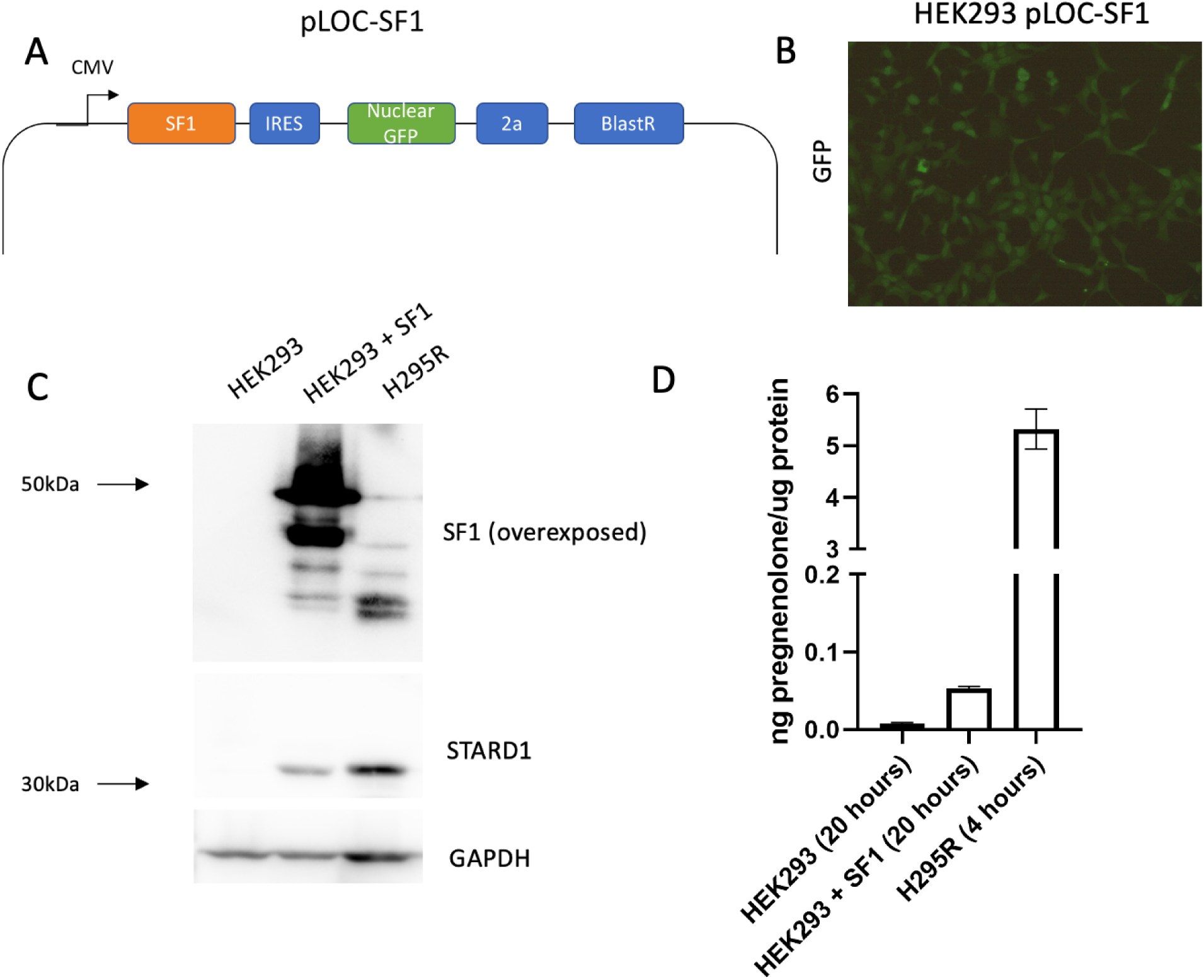
Constitutive expression of SF1 drives early steroidogenesis in HEK293 cells. **A:** Map of the pLOC-SF1 lentiviral vector for constitutive expression of SF1, a nuclear GFP marker and blasticidin resistance. The coding sequences are separated by an internal ribosome entry site (IRES) and 2a sequence. **B:** Image of GFP expression in HEK293 cells transduced with pLOC-SF1. **C:** Western blot comparing control HEK293 cells to transduced differentiated HEK293 cells. Steroidogenic H295R cells were compared as a positive control. **D:** Pregnenolone levels in media normalized to protein content of cells collected from the indicated cell lines. Media was incubated with cells for the indicated amount of time. Error bars represent the standard deviation of two technical replicates.

As a positive control we compared differentiated HEK293 cells with the H295R adrenocorticocarcinoma cell line. To facilitate comparison between different cell types, the total amount of pregnenolone in the media was normalized to the amount of protein in the well, obtained by lysing the cells and measuring protein concentration. Although steroidogenic HEK293 cells had much higher expression of SF1, there was lower expression of STARD1 and production of pregnenolone. Meanwhile, cortisol, the main glucocorticoid produced by adrenocortical cells, was not detectable. The fact that the differentiated HEK293 cells did not adopt a fully steroidogenic phenotype is not unexpected, as SF1 expression was unable to induce much steroidogenesis in more terminally differentiated cell types, such as HEK293 (13). Nevertheless, these findings confirm that lentiviral transduction of SF1 and steroidogenic differentiation approach works in our hands.

### iPSCs do not tolerate SF1 expression

Having confirmed that lentivirus could be used to effectively overexpress SF1 and differentiate HEK293 cells, we next tested transduction and differentiation directly on iPSCs. Transduced iPSCs showed little to no discernable GFP expression and most iPSCs were lost during the antibiotic selection process. When protein extract from the surviving cells were run on western blot, only a small amount of SF1 was detected (**Figure 2A**). A sample of media was collected from the cells to measure steroid levels (**Figure 2B**), however pregnenolone levels were not detected above background, and cortisol was not reliably detected at all in the treated iPSCs (data not shown).

**Figure 2:**
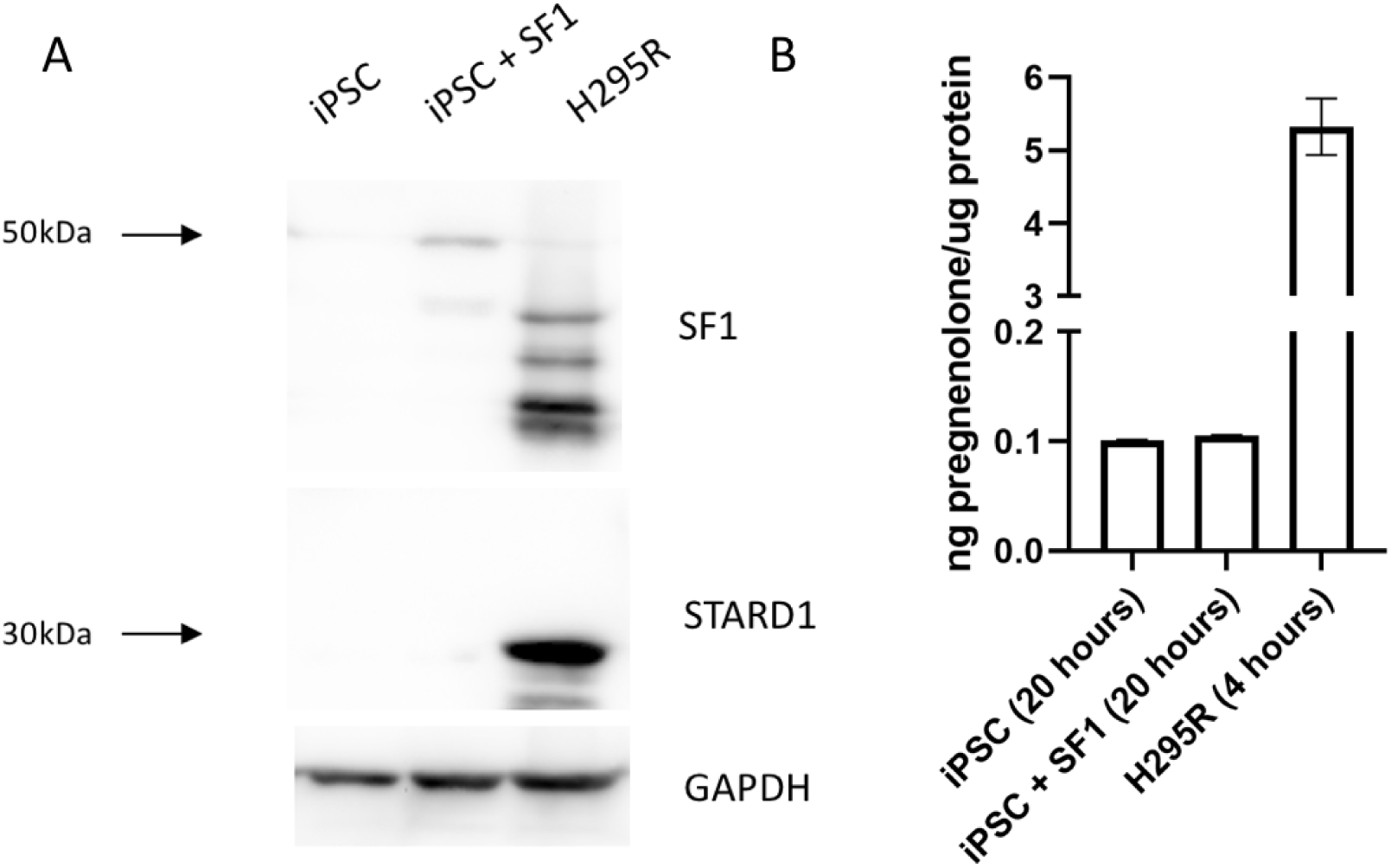
IPSCs do not tolerate SF1 overexpression. **A:** Western blot comparing iPSCs and transduced iPSCs to H295R cells as a positive control. **B:** Pregnenolone levels in media normalized to protein content of cells. Media was incubated with cells for the indicated amount of time. Error bars represent the standard deviation of two technical replicates.

### Differentiation of iPSCs to steroidogenic cells through an embryoid body intermediate

Previous literature shows that SF1 overexpression, specifically in pluripotent stem cells, (including iPSCs and embryonic stem cells) can be cytotoxic and ineffective at inducing steroidogenesis (11,13,23). One possible explanation is the ability of SF1 to dysregulate levels of pluripotency gene OCT4 (29). To tolerate SF1 overexpression, we adjusted our approach to partially differentiate the iPSCs so that they were no longer pluripotent, yet were still able retain sufficient developmental potential to become actively steroidogenic.

Existing approaches to differentiate pluripotent stem cells use a stepwise approach involving differentiation into an intermediate (8,20). Instead of costly commercial kits (20) or recombinant proteins (19), we tested simpler methods that use minimal additional reagents to differentiate iPSCs, prior to attempting SF1 lentiviral transduction. While some approaches led to cell death shortly after, or the cells did not produce detectable steroids following lentiviral transduction, we found that differentiating iPSCs into embryoid body (EB) intermediates, showed both cell survival and steroidogenesis **Figure 3 (Approach A)**. In comparison, an initial differentiation step using retinoic acid as described by Yazawa *et al.* (30) showed little evidence of steroidogenesis in our hands **Figure 3 (Approach B)**.

**Figure 3:**
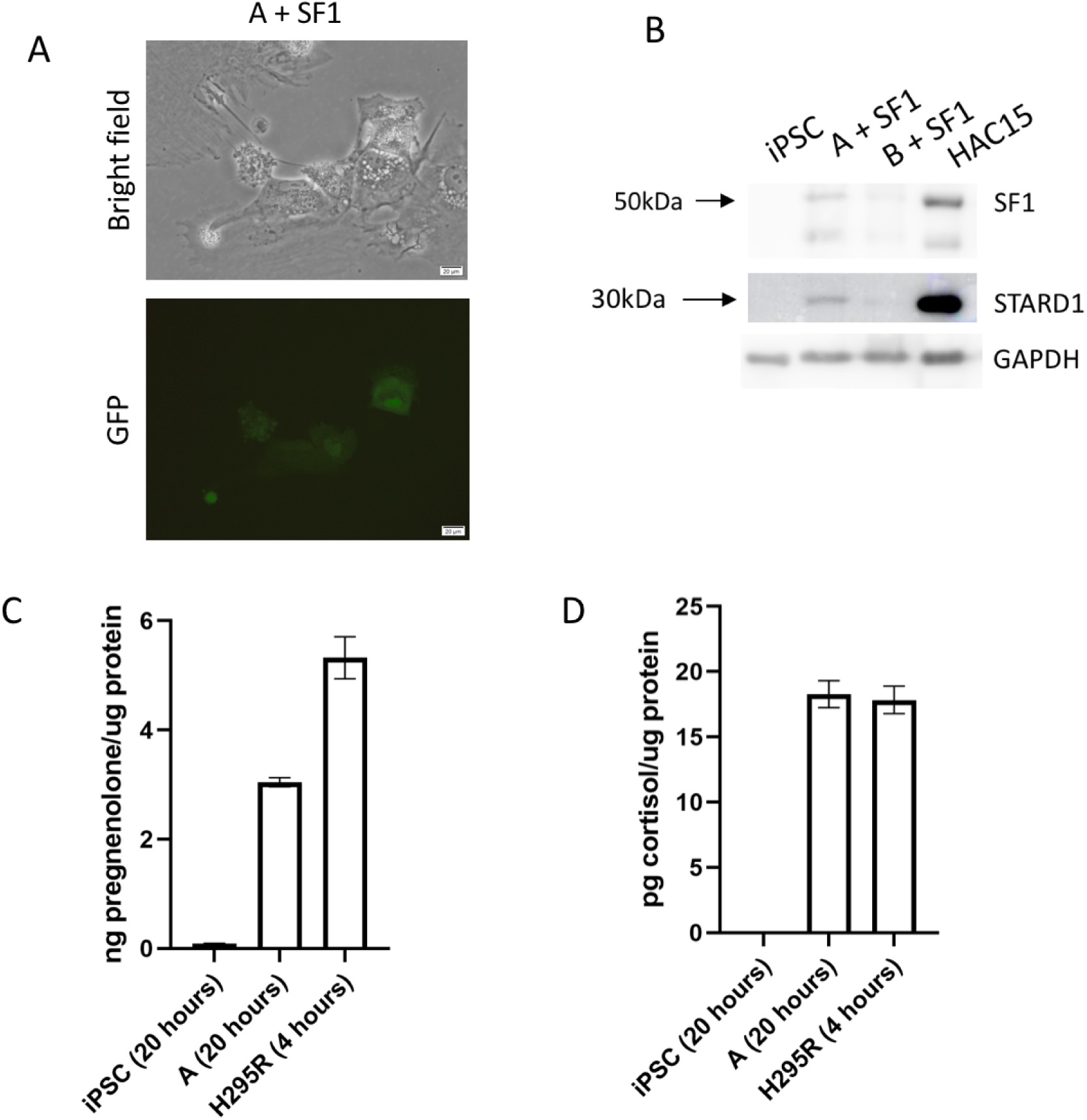
Partially differentiated iPSCs can tolerate SF1 overexpression and produce steroids including cortisol. **A:** Images of partially differentiated iPSCs transduced to express SF1 and nuclear GFP marker gene. **B:** Western blot comparing the starting iPSCs to two attempts at partially differentiated iPSCs transduced to express SF1 and STARD1, compared to steroidogenic HAC15 cells as a positive control. **C:** Pregnenolone and **D:** cortisol levels measured in media collected from cells for the indicated amount of time, and normalized to protein content of cells. Media from H295R cells was collected as a positive control. Error bars represent the standard deviation of two technical replicates.

Differentiating iPSCs into an EBs, which are three-dimensional floating aggregates of cells, allows stem cells to be further differentiated into germ layers (27). First, EBs were formed (**Figure 4A**) by passaging cells into a non-tissue culture-treated plates (e.g. a sterile Petri dish). Next, EBs were plated into tissue-culture treated plates, where EB intermediate outgrowing cells were observed as adherent cells grew from the cell aggregates. Notably, these EB intermediate cells had reduced expression of OCT4, a pluripotency gene, and increased expression of brachyury, an early mesoderm marker (**Figure 4B**). Together, these findings suggest the EB intermediates were no longer pluripotent and had potentially differentiated into germ layer cells.

**Figure 4:**
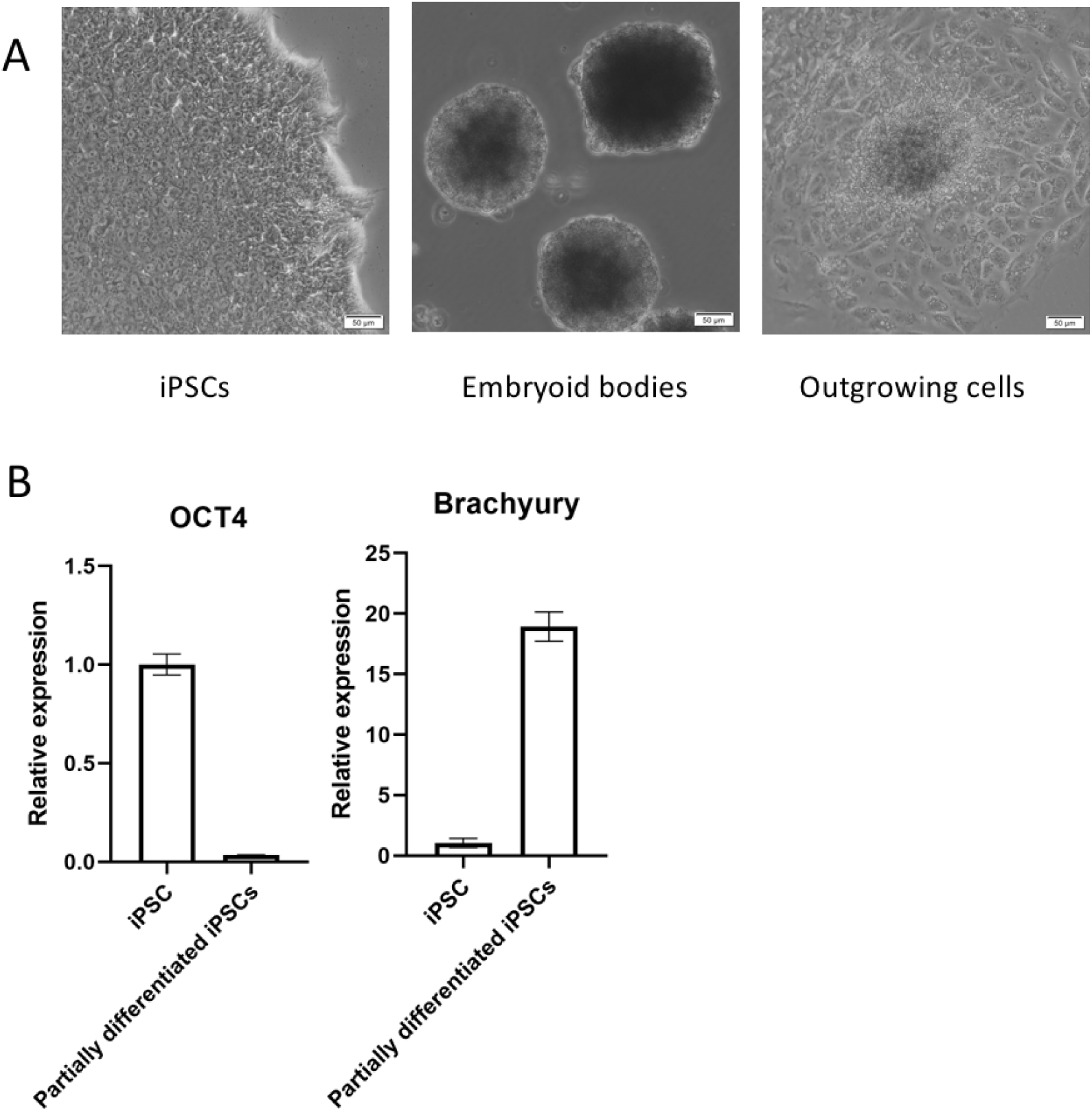
Partially differentiated iPSCs through an embryoid body intermediate. **A:** Images of the process of partially differentiating iPSCs via embryoid body formation. **B:** RT-qPCR of pluripotency marker OCT4 and mesoderm marker brachyury compared between the starting iPSCs and the embryoid body differentiated cells. Error bars represent the standard deviation of three technical replicates.

With respect to differentiation into a steroidogenic state, the EB intermediates transduced with pLOC-SF1 expressed the nuclear GFP marker. Moreover, differentiated cells expressed similar levels of SF1 to steroidogenic adrenocortical cells (**Figure 3B**). Compared to HEK293 cells, where very high SF1 overexpression was needed to see slight induction of STARD1 (**Figure 1C**), moderate SF1 expression was able to induce more comparable levels of STARD1 to steroidogenic cells (**Figure 3B**). In addition, the differentiated cells synthesized detectable levels of pregnenolone that were well above background (**Figure 3C**). Critically, these cells were also capable of producing detectable amounts of cortisol (**Figure 3D**). When normalized to protein content, these cells were comparable to, but still less steroidogenic, than adrenocortical cells. Taken together, these findings show that following initial differentiation into EB intermediates, SF1 lentiviral transduction and steroidogenic differentiation was able to activate all the genes needed for glucocorticoid synthesis.

### Further optimization and characterization of the differentiated cells

Notably, we observed that the time point at which EB intermediates are transduced with SF1 following plating impacts the differentiation into a steroidogenic state. Assessing brachyury expression from EB-differentiated iPSCs harvested at different time points showed brachyury expression declined over time (**Figure 5A**), suggesting the partially differentiated cells continue to change following EB formation and may only be amenable to differentiation at certain time points. To determine the ideal time for differentiation, we tested lentiviral transduction of the same set of differentiated iPSCs at 4, 7 and 10 days after plating of the embryoid bodies. Cells transduced 7 days post plating showed the greatest SF1 lentiviral transduction level, STARD1 expression and steroid levels (**Figure 5B-D**), suggesting that this was the ideal timepoint.

**Figure 5:**
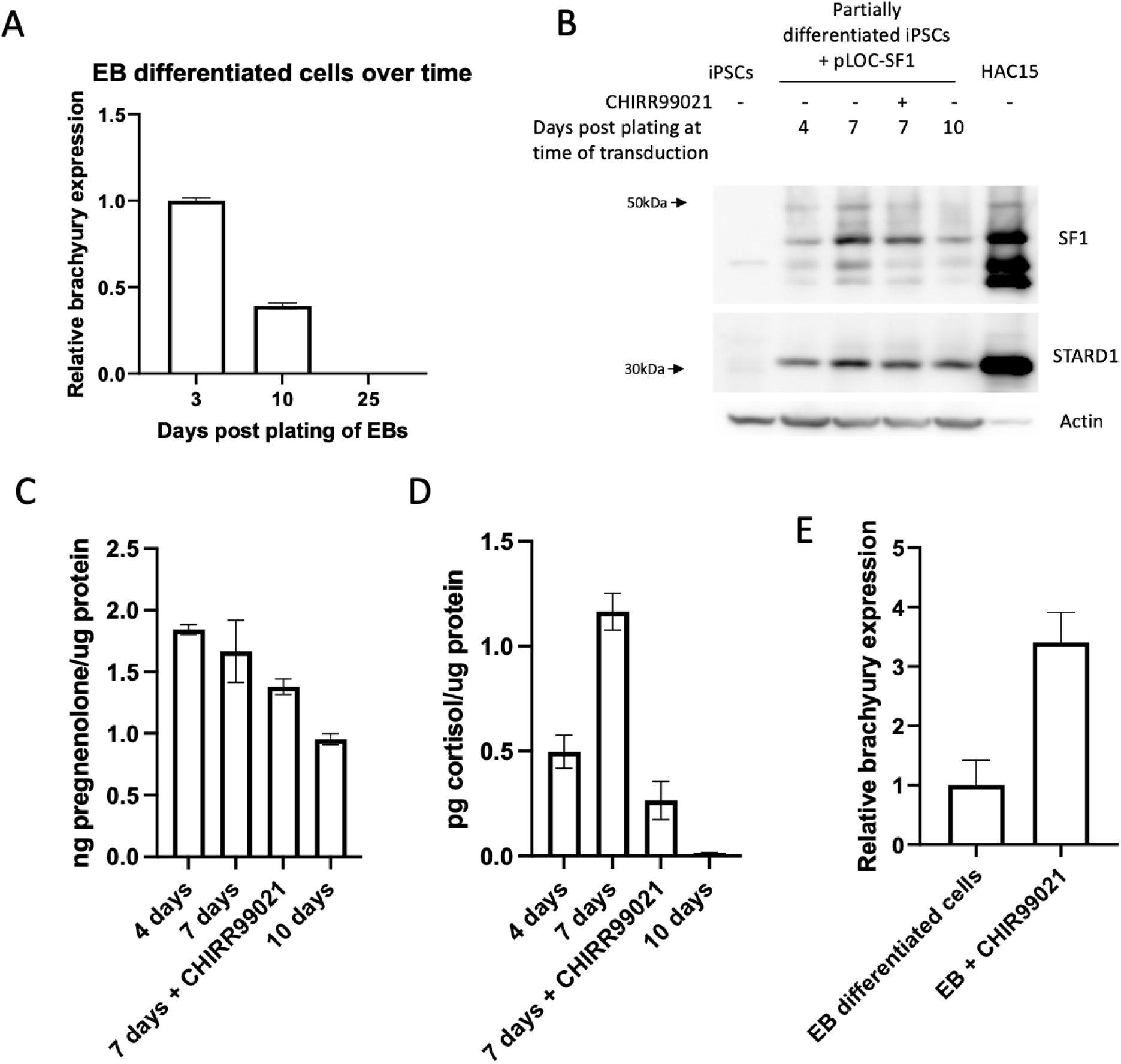
Further optimization of differentiation of iPSCs to steroidogenic cells. **A:** Brachyury expression from EB-differentiated cells collected at four time points after plating of the embryoid bodies. Error bars represent standard deviation of three technical replicates. **B:** Western blot of partially differentiated iPSCs transduced at either 4, 7, or 10 days post plating of the embryoid bodies. One set of cells was also treated with 3 μM CHIR99021 from 24-72 hours of embryoid body formation process, and then transfected at 7 days post plating of the embryoid bodies. HAC15 cells were used as positive control. **C:** Pregnenolone and **D:** cortisol levels measured in media collected from the transduced cells overnight. Values were normalized to the protein content of each sample to account for any differences in cell density. Each sample is labelled by the number of days post plating of the EBs when they were transduced (4-10 days). Error bars represent the standard deviation of two technical replicates. **E:** Expression of brachyury detected using RT-qPCR from EB differentiated cells treated with CHIR99021 or DMSO from 24-72 hours of embryoid body formation process. Cells were collected for qRT-PCR 3 days after plating the embryoid bodies.

We were also curious whether small molecules that can promote mesodermal development might enhance differentiation into steroidogenic cells, as the adrenal cortex originates from the intermediate mesoderm (22), and as previous work generated steroidogenic cells from pluripotent stem cells via a mesodermal intermediate (18). Thus, we tested if we could improve steroidogenic differentiation with the Wnt agonist CHIR99021, which increases in brachyury expression and enhances mesodermal differentiation of stem cells (31,32). We confirmed that treatment with CHIR99021 during embryoid body growth increased brachyury expression (**Figure 5E**). However, this treatment did not increase SF1 or STARD1 expression (**Figure 5B**), pregnenolone production (**Figure 5C**) or cortisol production (**Figure 5D**), suggesting elevated brachyury expression in partially differentiated cells did not improve the capacity for steroidogenic differentiation. Based on these findings, the final workflow for subsequent generation of cortisol-producing cells is detailed in **Figure 6**. A detailed stepwise description of the iPSC differentiation protocol can be found in the **Supplemental Methods**.

**Figure 6:**
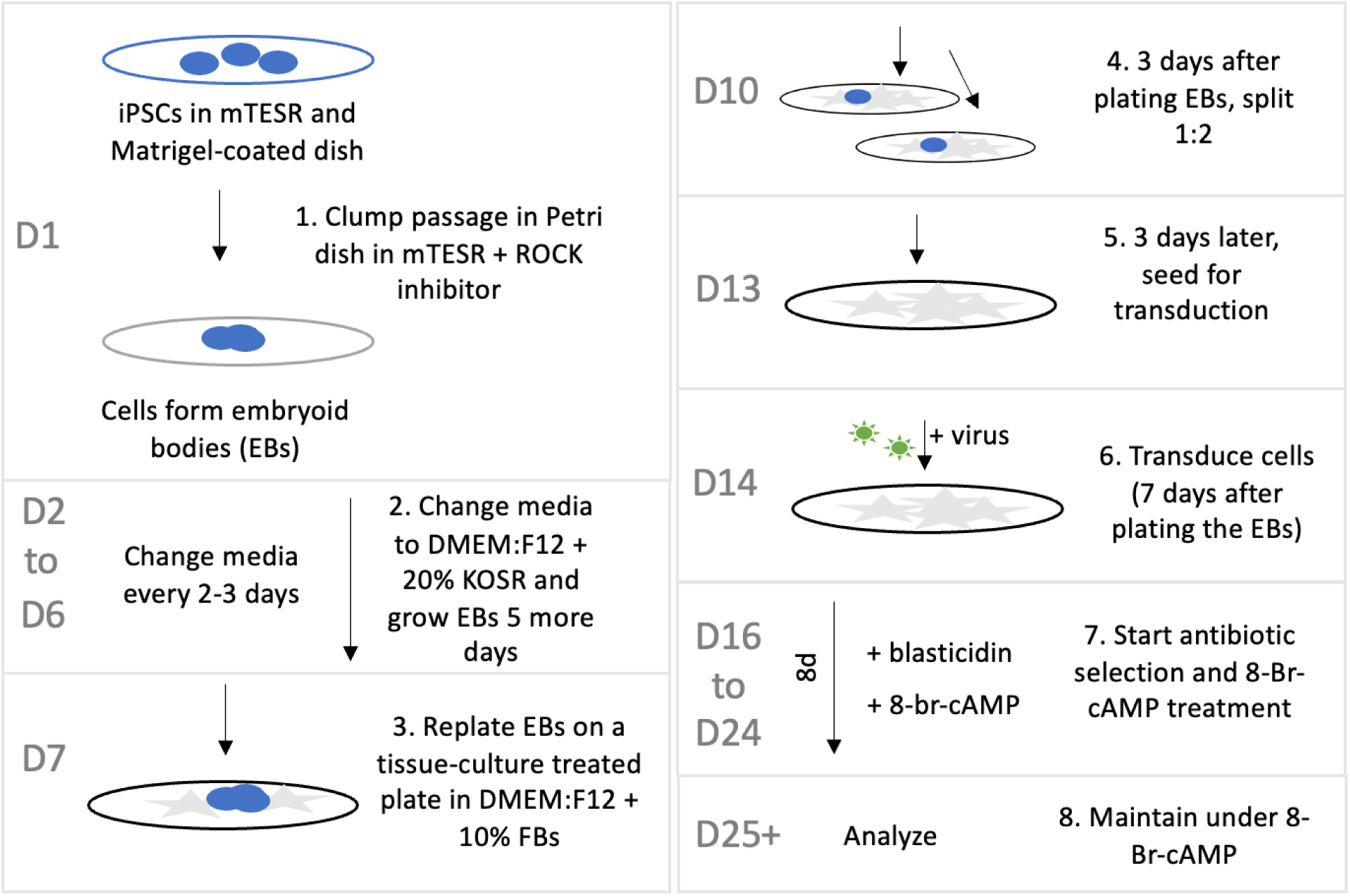
Overall workflow for generating SF1 expressing cells. Flowchart illustrating the general protocol for differentiating iPSCs into steroidogenic cells starting by inducing embryoid bodies and subsequent outgrowth, followed by transduction with SF1 lentivirus, and treatment with 8-Br-cAMP and blasticidin selection. Detailed steps for the differentiation protocol can be found in the Supplemental Methods.

### Reproducibility and long-term stability

To test the consistency of the protocol, three plates of iPSCs were differentiated in parallel and steroid levels were assessed two weeks post transduction (**Figure 7A-B**). Cells were also kept in culture for 1-month post-transduction. As noted in other protocols generating steroidogenic cells (7,18,21,33), the differentiated cells did not appear to proliferate. Rather, the cells also gradually deteriorated, with over half of the cells lost by the end of the month.

**Figure 7:**
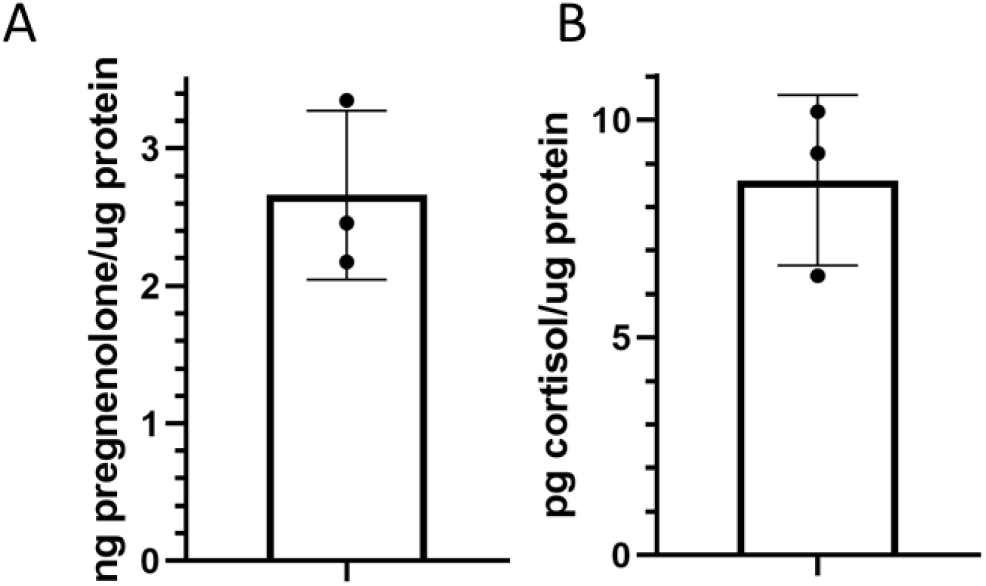
Assessing consistency of final protocol. Three plates of iPSCs were differentiated according to the protocol in parallel. Media was collected from the cells and steroid levels were measured and normalized to protein content of each sample. Error bars reflect the standard deviation of the three parallel replicates.

## Discussion

We describe a novel and simple approach to differentiate iPSCs into a steroidogenic state that combines an embryoid body differentiation step with lentiviral SF1 overexpression and cAMP treatment. We confirmed that the differentiated cells express STARD1, a key mediator of steroidogenesis, and produce detectable levels of both pregnenolone (the first steroid precursor) and cortisol (a downstream steroid hormone). While this approach will facilitate studies investigating steroidogenesis in patient-derived iPSCs, further characterization of the properties of these iPSC-derived steroidogenic cells would be beneficial and there are future optimizations that could be developed. Additionally, certain limitations of the protocol should be considered.

To better understand these iPSC-derived steroidogenic cells, it would be beneficial to characterize the expression levels of other steroidogenic mediators including key enzymes and the ACTH receptor, as well as look at the production of additional steroid hormones. In this context, assessing the levels of Leydig-specific steroidogenic enzymes and hormones would help determine if cells are directed towards androgen synthesis. Further optimization could be performed to ensure differentiation is directed towards adrenocortical or Leydig-like cells (20).

One potential limitation is experimental variability, which may limit comparisons of subtle differences in steroid production between cell lines. For example, similar to the approach we report here, Ishida *et al.* used an EB differentiation and doxycycline-induced SF1 expression approach to promote Leydig cell differentiation, and found resultant testosterone production varied over a 4-fold range between different experiments (8). In this regard, our findings show the importance of timing following the EB differentiation step, which may contribute to inconsistency. While the significance of brachyury as a phenotypic marker is unclear, it may affect the cells’ propensity for differentiation. An alternative and more consistent initial differentiation step could improve reproducibility.

Another potential limitation is that the iPSC-derived differentiated steroidogenic cells do not proliferate and also degenerate in prolonged culture, limiting their use for longer-term experiments. Limited lifespan of steroidogenic cells has also been observed in other studies where cells were reported to survive 25-28 days before showing degeneration (8,21). Further refinement of cell culture conditions may be able to extend the lifespan of these cells.

Overall, expanding upon the lentivirus-based transduction protocol to generate adrenocortical-like cells initially established by Ruiz-Babot *et al.* (7), we were able to extend this method to iPSCs by introducing an additional embryoid body differentiation step. The production of cortisol-producing cells from iPSCs uses a straightforward protocol that does not require specialized reagents, thus lowering costs and the required expertise.

## Acknowledgements

This work was supported by grants from the Calgary Parkinson Research Initiative through the Hotchkiss Brain Institute (TES) and the Canadian Institutes of Health Research (TES). LLG was supported by a Cumming School of Medicine Graduate Scholarship.

## Author contributions

LPL-G designed, performed, and analyzed experiments, prepared figures, and wrote and revised the manuscript. TES conceived and designed the study, analyzed experiments, supervised the study, and wrote and revised the manuscript.

## Supplemental Methods

Detailed protocol for steroidogenic iPSC differentiation

Reagents:

- Culture reagents for iPSCs: mTESR Plus (Stemcell, 100-0276) on Matrigel (Corning, 354277) coated plates
- Media for embryoid bodies: DMEM-F12 with 20% Knockout Serum Replacer (KOSR) (Gibco, 10828010)
- Media for replated partially differentiated cells: DMEM-F12 with 10% heat-inactivated fetal bovine serum
- Y-27632 ROCK inhibitor (Stemcell 72308), use at final concentration of 10uM
- Polybrene (Sigma-Aldrich, 107689), use at final concentration of 10ug/mL
- Blasticidin (Invivogen, ant-bl), use at final concentration of 10ug/mL
- 8-bromo-cAMP (Stemcell 73604), use at final concentration of 100uM

Procedure:

- Seed a Matrigel-coated 35mm dish with 2×10^5^ iPSCs (fairly low confluency to allow time for colonies to grow) in mTESR Plus and on a Matrigel-coated dish.
- Day 1: Around 4-5 days later, once plate is about 70% confluent, clump passage iPSCs using EDTA dissociation into 4mL of mTESR + ROCK inhibitor. Transfer cells to a 60mm non-tissue culture-treated plate, such as a sterile Petri dish.
  - To maximize cell survival and embryoid body (EB) formation, ensure cells are not overly confluent. We found that passaging cells directly into DMEM:F12 + 20% KOSR reduces cell survival and EB formation. In the absence of ROCK inhibitor, cell survival and EB formation is also reduced.
- Day 2: Change media to DMEM:F12 + 20% KOSR.
  - Collect the media and any floating EBs into a 15mL tube and let settle about 10 minutes. Remove the supernatant and gently resuspend EBs (do not dissociate) in the new media.
- Day 3-6: Culture in DMEM:F12 with 20% KOSR, changing media every 1-2 days.
  - After the first couple of days, the media can acidify quickly and will need to be changed every day. Change the media as described above.
- Day 7: After growing the EBs for 6 days, plate in DMEM:F12 with 10% FBS in a tissue culture-treated 60mm dish.
  - Collect the media and EBs, let settle 10 min, resuspend in 4mL DMEM:F12 with 10% FBS and plate in a tissue culture-treated plate. From here on out, continue to grow the cells on tissue-cultured treated plates.
- Day 10: After 3 days (or sooner, if the plate is confluent), split the plate into either two 60mm plates or one 10cm plate.
  - Passage the cells with Accutase or trypsin. Include ROCK inhibitor in the media to minimize cell death.
- Day 13: After 3 more days (6 days post plating of the embryoid bodies), seed for transduction at a density of 3-4.5×10^4^ cells per cm^2^
  - Seeding cells for transduction at this timepoint is usually enough time for the cells proliferate, while also remaining within the window during which they can be effectively differentiated.
  - Seed an additional plate of cells to use as an antibiotic control to determine when antibiotic selection is complete.
- Day 14: Transduce the cells by adding 400 viral particles/cell of pLOC-SF1 virus and 10ug/mL polybrene.
  - As an alternative to measuring physical titre, we found good transduction efficiency using virus collected from a dish three times the size (i.e. for a 10cm dish of cells, transduce with virus collected from 3 10cm dishes of cells, or collected 3 times).
- Day 15: Change media
- Day 16: Two days post transduction, start antibiotic selection using 10ug/mL blasticidin and treatment with 100uM 8-Br-cAMP.
  - Refresh the antibiotics and cAMP every 2-3 days.
  - Treat the control plate of untransduced cells in parallel to determine when antibiotic selection is complete (around 7 days).
  - Treat cells with cAMP for at least 8 days before analysing cells.
- Day 25: After 8 days of 8-Br-cAMP treatment, the media or cells can now be collected for analysis. Maintain cells under 8-Br-cAMP for continued culture.

